# A mismatch between human early visual cortex and perception in spatial extent representation: Radial bias shapes cortical representation while co-axial bias shapes perception

**DOI:** 10.1101/2023.02.16.528416

**Authors:** Juhyoung Ryu, Sang-Hun Lee

## Abstract

An object occupies an enclosed region in the visual field, which defines its spatial extent. Humans display exquisite finesse in spatial extent perception. Recent series of human neuroimaging and monkey single-cell studies suggest the spatial representation encoded in the early visual cortex (EVC) as the neural substrate of spatial extent estimation. Guided by this “EVC hypothesis” on spatial extent estimation, we predicted that human estimation of spatial extents would reflect the topographic biases known to exist in EVC’s spatial representation, the co-axial and radial biases. To test this prediction, we concurrently assessed those two spatial biases in both EVC’s and perceptual spatial representations by probing the anisotropy of EVC’s population receptive fields, on the one hand, and that of humans’ spatial extent estimation, on the other hand. To our surprise, we found a marked topographic mismatch between EVC’s and perceptual representations of oriented visual patterns, the radial bias in the former and the co-axial bias in the latter. Amid this topographic mismatch, the extent to which the anisotropy of spatial extents is modulated by stimulus orientation is correlated across individuals between EVC and perception. Our findings seem to require a revision of the current understanding of EVC’s functional architecture and contribution to visual perception: EVC’s spatial representation (i) is governed by the radial bias but only weakly modulated by the co-axial bias, and (ii) do contribute to spatial extent perception, but in a limited way where additional neural mechanisms are called in to counteract the radial bias in EVC.

**Significant statement:** Previous anatomical and functional studies suggest both radial and co-axial biases as topographic factors governing the spatial representation of the early visual cortex (EVC). On the other hand, EVC’s fine-grained spatial representation has been considered the most plausible neural substrate for exquisite human perception of spatial extents. Based on these suggestions, we reasoned that these two topographic biases are likely to be shared between EVC’s and perceptual representations of spatial extents. However, our neuroimaging and psychophysics experiments implicate a need for revising those two suggestions. Firstly, the co-axial bias seems to exert only a modulatory influence on EVC’s functional architecture. Secondly, human spatial extent perception requires further contribution from neural mechanisms that correct EVC’s spatial representation for its radial bias.

## Introduction

An object occupies an enclosed region in the visual field, which defines its *spatial extent*. For primates, who skillfully manipulate objects with hands, accurately estimating objects’ spatial extents is crucial for successful adaptation to their surroundings (Gibson, 1966). For instance, to pick up the eggs differing subtly in shape, dexterous finger manipulation must be supported by accurate spatial extent estimation (Horton et al., 2012). Attesting to the importance of spatial extent estimation, humans can discriminate subtle differences between objects’ aspect ratios with exquisite finesse (Regan and Hamstra, 1992; Regan et al., 1996).

Given this finesse of estimation performance and the topographic nature of spatial extents, the high-resolution spatial representation in the early visual cortex (EVC, V1/2/3) has been favored as a neural substrate of spatial extent estimation (Michel et al., 2013; Schwarzkopf, 2015). This EVC hypothesis gained empirical support from a series of human neuroimaging (Murray et al., 2006; Fang et al., 2008; Sperandio et al., 2012; Pooresmaeili et al., 2013; He et al., 2015; Moutsiana et al., 2016; Ho and Schwarzkopf, 2022) and monkey single-cell (Ni et al., 2014) studies, where EVC’s spatial representations were contextually modulated in the way matching perceptual representations.

According to the EVC hypothesis, human perception of spatial extents is likely subject to the biases known to govern the spatial representations in EVC or its preceding neural stages. Two such biases are particularly pertinent to the neural encoding of spatial extents: *co-axial bias* and *radial bias*. Here, *bias* refers to any topographic properties of neurons or their connections that lead to anisotropic representations of objects’ spatial extents in visual space, co-axial when elongated along the axis aligned to an object’s orientation and radial when elongated along the radial axis on which the object is positioned. These biases have been reported at multiple levels, including the shape of single or aggregated cells’ receptive fields (RFs and pRFs, respectively) and horizontal connections between cells. The co-axially biased spatial extents were found in EVC’s RFs (Jones and Palmer, 1987; Anzai et al., 1999) and long-range horizontal connections (Bosking et al., 1997; Sincich and Blasdel, 2001; Iacaruso et al., 2017). On the other hand, the radially biased spatial extents were found in the RFs of animals’ retinal ganglion cells (Leventhal and Schall, 1983; Schall et al., 1986a, 1986b; Watanabe and Rodieck, 1989; Passaglia et al., 2002) and humans’ RGCs (Rodieck et al., 1985), and in the pRFs of human V1 (Merkel et al., 2018, 2020).

These two lines of evidence appear to indicate that human EVC’s spatial representation is likely governed by both biases. However, the correlation between the preferred orientations and polar-angle positions of single cells (Leventhal, 1983; Leventhal et al., 1984; Bauer et al., 1989; Durand et al., 2007) or cortical sites (Freeman et al., 2011, 2013; Sun et al., 2013; Fang et al., 2022; Roth et al., 2022) allows the possibility that one of the biases is absent or negligible but only appears to exist due to its shared variability with the other genuine bias. Furthermore, the two biases have been reported in a variety of species, which adds uncertainty to whether the same biases exist in human EVC. Thus, three scenarios are left unresolved: only the co-axial bias (Fig. 1A), only the radial bias (Fig. 1B), or both regulate(s) the spatial extent representation in human EVC.

**Figure 1.**
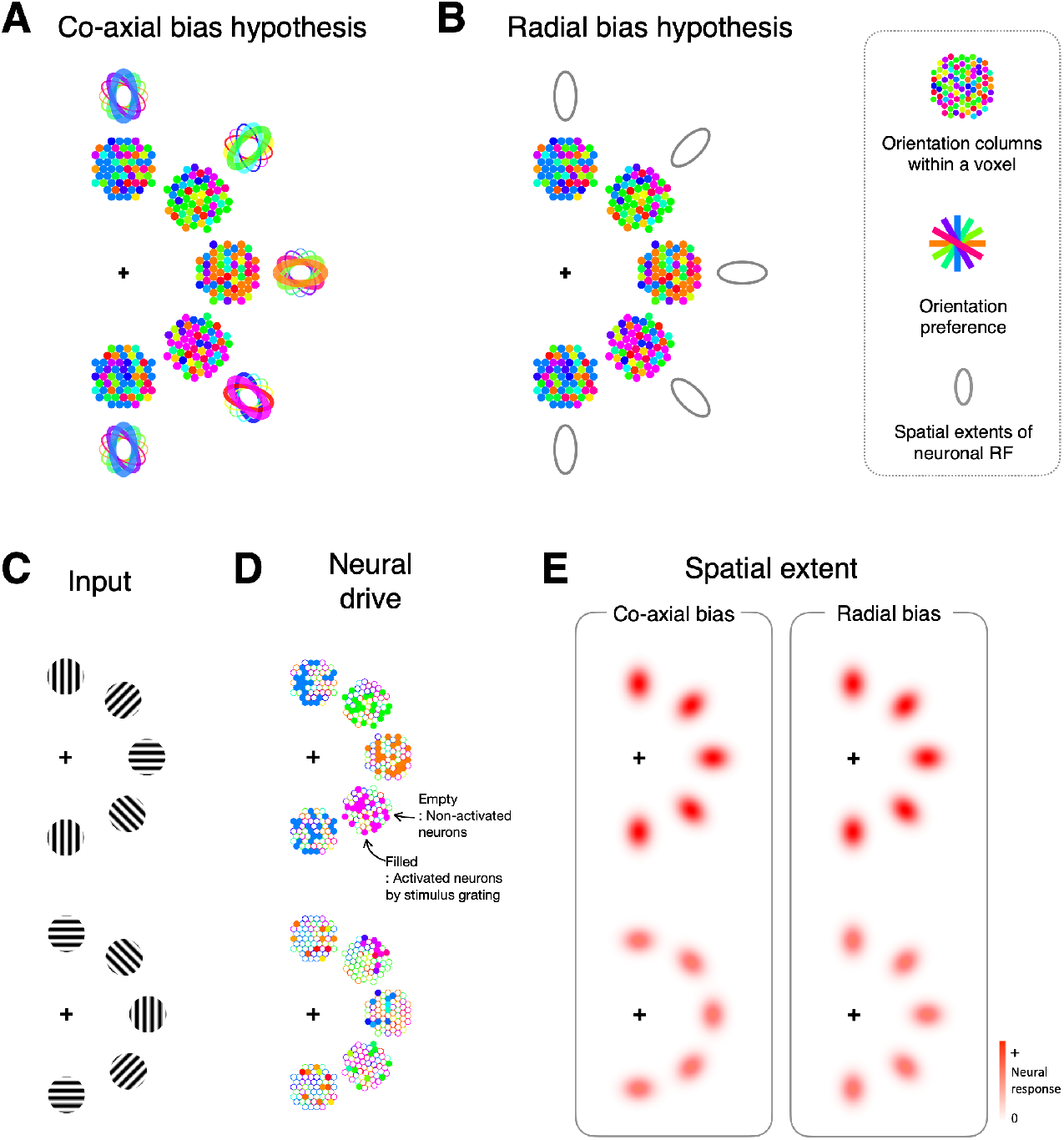
Rationale for assessing the spatial bias in pRF using oriented patterns. ***A-B***, Schematic illustrations of the co-axial and radial (***B***) bias hypotheses about the topographical factor governing the spatial representation of EVC. The fixation location is marked by crosshairs. Each cluster of colored dots represents an aggregated orientation columns within a voxel representing a given retinotopic position. The colors represent orientation preference, and their distributional biases reflect the strong correlation between columns’ preferred orientation and polar-angle position. The ellipsoids surrounding the dot clusters represent the topographical bias of the orientation column. Note that the direction of topological bias in any given orientation column is determined by the axis of its preferred orientation according to the co-axial bias (***A***) and by the radial axis to which the voxel’s retinotopic position belongs (***B***). ***C-E***, Radially and tangentially oriented gratings (***C***), the neural drives by those gratings (***D***), and the spatial extents evoked by those neural drives (***E***). The oriented gratings (***C***) will provide the selective drive to the columns within single voxels (***D***). Then, the spatial extents will be determined by the orientation columns that received the selective drive (***E***). As a result, due to their difference in what it is the factor determining the topographical biases (***A***,***B***), the two hypotheses predict the tangentially oriented gratings to result in the spatial extents that are elongated in the different axes in EVC’s spatial representation.

To resolve this issue, we created a critical condition where the two biases make conflicting predictions about the anisotropy of spatial extents (Fig. 1C-E) and evaluated which scenario best explains pRFs’ spatial extents using functional magnetic resonance imaging (fMRI). Having found that the radial bias predominates over the co-axial bias in EVC’s spatial representation, we verified the EVC hypothesis by checking whether the same radial bias regulates perceived spatial extents. To our surprise, the co-axial bias regulated perceived spatial extents, which are at odds with the EVC hypothesis and call for the reinterpretation of previous findings.

## Materials and Methods

### Subjects

Twenty-nine human subjects (normal or corrected-to-normal vision; aged 20-30; 15 females) participated in the fMRI experiment. Twenty-seven of them (14 females) also participated in the psychophysical experiment (14 females). All subjects except for one (the first author) were naïve to the purpose of the study. They provided informed consent, and the experiments were conducted by the guidelines approved by the Institutional Review Board at Seoul National University.

### Methods for FMRI experiment

#### MRI acquisition

MR data were acquired using a 3T Siemens MAGNETOM Trio at the Seoul National University Brain Imaging Center. For anatomical image scans, a high-resolution T1-w MPRAGE was acquired using a 32-channel head coil (1 × 1 × 1 mm^3^ voxel, 1.9s TR, 2.36 ms TE, 9° flip angle (FA) for 12 subjects; 0.85 × 0.85 ×0.85 mm^3^ voxels, 2.4s TR, 3.42 ms TE, 8° FA for 17 subjects). A 20-channel head coil (only the bottom part of a 32-channel head coil) was used for functional image scans to avoid partial blockage of the visual field. Twenty-four oblique slices orthogonal to the Calcarine sulcus were prescribed at the most posterior part at the occipital pole. GRAPPA, 1.5 s TR, 30 ms TE, 75° FA, 2 × 2 × 2 mm^3^ voxel size, 96 × 80 matrix size, two acceleration factors, and the interleaved slice acquisition order with 62.5 ms inter-slice interval were used. Functional scan follows in-plane T1-w imaging using a 20-channel head coil (MPRAGE, 1 × 1 ×1 mm^3^ voxel, 1.6 s TR, 2.36 ms TE, 9° FA). It is to compensate head motion between 32-channel T1-w scan and functional scan.

#### Visual stimulus presentation

Stimuli were generated using MATLAB (MathWorks) in conjunction with PsychToolbox (Brainard, 1997) on a Macintosh computer. The stimuli were presented by an LCD projector (Canon XEED SX60; Canon) at its native resolution (1,400 × 1,050 pixels; refresh rate, 60 Hz) onto a rear projection screen placed inside the magnet bore. The distance to the screen from the eyes was 87 cm, and the projection area on the screen was 34.5 cm × 26 cm, resulting in a visual angle of 22° (width) × 17° (height). Subjects viewed the stimuli through the front surface of a mirror with a multilayer dielectric reflective coating (Sigma Koki) that was mounted on the head coil. A custom-made neutral density filter (9% transmission rate; Taeyoung Optics) was inserted between the projector lens and the screen to control the overall level of stimulus luminance. The color lookup table was calibrated to linearize the luminance values at the screen center ranging from 0.0045 cd/m^2^ to 63.5 cd/m^2^ by using a luminance meter (LS-100; Konica Minolta Sensing in conjunction with in-house software for automated measurement and correction.

#### Visual stimulus design

To assess whether the spatial extents of pRF of the V1, V2, and V3 voxels are anisotropic in the polar space, and, if so, how the direction of the strength of such anisotropy depends on stimulus orientation or the polar angle position of the pRF, we acquired fMRI data from the occipital lobes of 29 human subjects while they were viewing radially or tangentially oriented narrowband orientation stimuli drifting slowly around a fixation point. Specifically, instead of counter-phased flickering checkerboards in conventional methods (Dumoulin and Wandell, 2008), multiple orientation carriers traverse the visual field (Park et al., 2013a): high-contrast and counter-phased flickering (1/150 ms) Gabor patches drift across the visual field to evoke cortical activities as it crosses over an aggregated RFs of neurons within a given cortical tissue. Since we were interested in the anisotropic spatial extent of pRF over the radial versus tangential axes of the polar space depending on stimulus orientation, we considered two orthogonal orientation conditions defined in terms of polar coordinate system: We used Gabor patches with radial or tangential orientation arranged in the shape of ring or bowtie wedge (See the first and the second row in Fig. 2B; Radial orientation condition is shown for example), and ring-or wedge-shaped aperture traverses the visual field in the radial or tangential direction. The aperture follows a periodic pattern and completes a cycle in 24 s, and the 24 s periodic pattern repeats nine times with no interval between cycles. In each cycle, the aperture moved in sync with the fMRI data acquisition (steps in every 1.5 s), and flickering Gabor stimuli in 1/150 ms remained at a given position for 300 ms, and a stimulus-off period of 75 ms was followed (For schematics, see Fig. 2A). The stimuli covered spatially extended region of retinotopic space up to 7.9° in radius.

**Figure 2.**
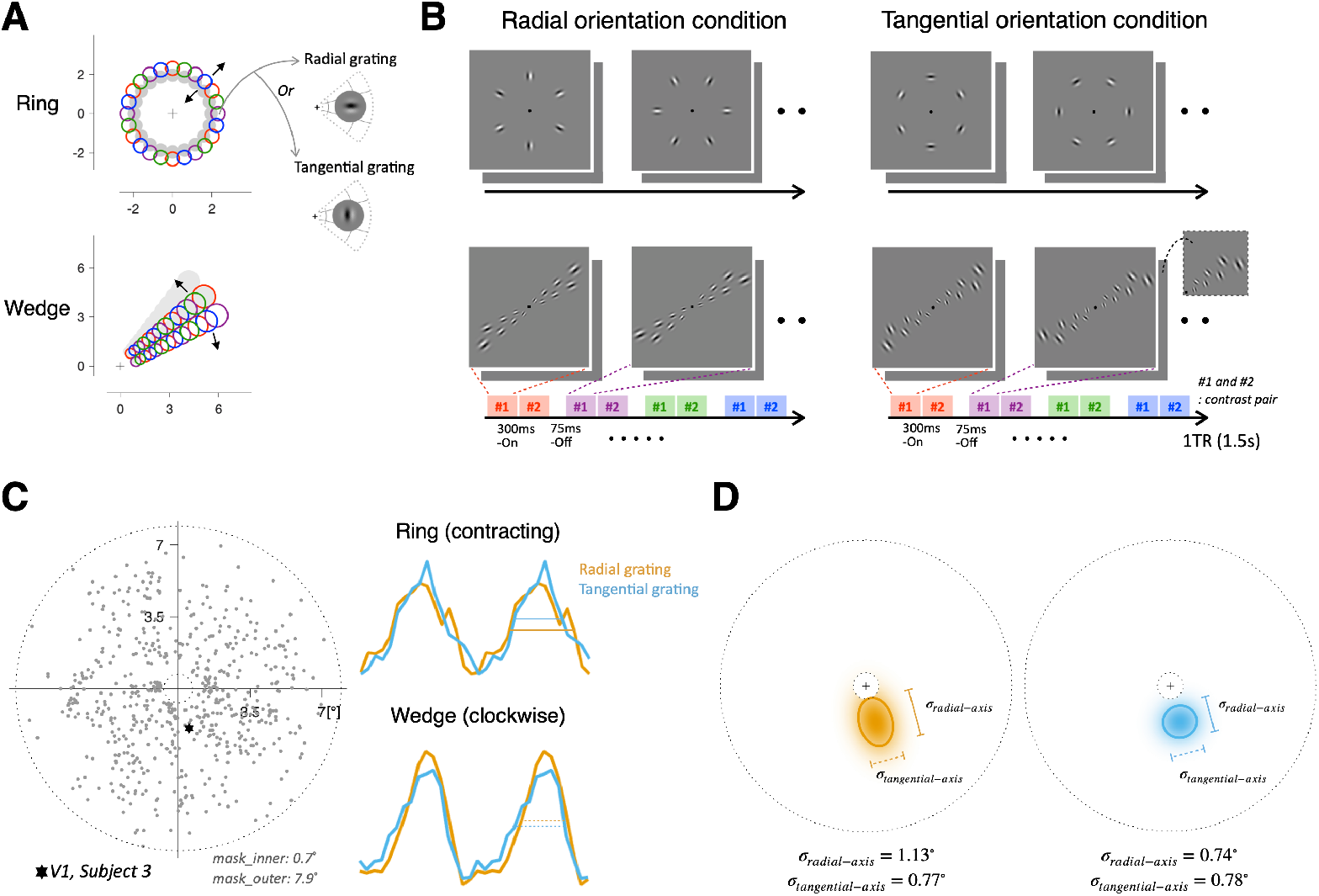
FMRI experimental design for estimating pRFs. ***A-B***, Spatial arrangements (***A***) and temporal sequence **(*B*)** of pRF mapping stimuli. At any given ring-shape or wedge-shape aperture, Gabor stimuli appeared only within one of the four packs of apertures (circles with the same color, one color for one pack, shown in ***A***) in the fixed order (specified at the bottom of ***B***). The ring-shape aperture traversed the visual field in the radial direction, either expanding or contracting (black arrows in ***A***), whereas the wedge-shape aperture in the tangential direction, either clockwise or counter-clockwise). Each aperture remains at a given position for 1.5 s (1TR) and shifts by the size of the half-width of the aperture (Gray shaded area in ***A***) to make sure that the stimulus was turned on for 3 s at any given visual space. Within each pack of Gabor stimuli, we did not allow for spatial overlap between neighboring Gabor stimuli to minimize the second-order interaction effects. ***C-D***, Estimation of pRF positions (***C***) and spatial extents (***D***). For single voxels, we estimated their pRF centers (***C***, left) and the spatial extents (***D***) by fitting the 2D Gaussian function to their time courses of fMRI responses voxels to the stimuli illustrated in ***B*** (***C***, right). The time courses of fMRI responses (***C***, right) came from an example voxel whose pRF center is marked by the star in the visual field (***C***, left) and whose pRF spatial extents are shown in ***D***. The colors of the fMRI time series and pRF spatial extents represent the stimulus orientation conditions.

To provide an effective trigger at any visual field, first, we make sure that visual stimuli stay 3 s (2TR) in the whole visual field: ring-or wedge-shaped aperture remained at a given position for 1.5 s (1TR) and is followed by the next stimulus aperture in a new position, that was shifted by the size of the half-width of the aperture. Second, the Gabor size increases linearly as a function of the eccentricity position of the Gabor center, and spatial frequency values were determined by their eccentricity position by applying the following logarithmic function: 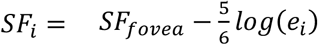. *SF*_*i*_ is the spatial frequency of Gabor located in the eccentricity of *i*, and *SF*_*fovea*_ is the spatial frequency at the fovea as 3 cycles/°. As a result, the spatial frequency ranged from 1.34 cycle/° at an eccentricity of 7.3° to 3 cycle/° at an eccentricity of 1°.

To minimize potential second-order interaction effects between neighboring Gabor patches, e.g., collinear facilitation, we imposed constraints on the spatiotemporal layout of the contrast envelops of Gabor stimuli, and we randomly varied the phase of Gabor stimulus. Details for ring and wedge scans are below, respectively.

For the expanding/contracting ring scan, 16 ring apertures of different sizes centered at eccentricity positions, [1°, 1.30°, 1.63°, 1.95°, 2.31°, 2.67°, 3.05°, 3.44°, 3.86°, 4.29°, 4.75°, 5.21°, 5.71°, 6.21°, 6.76°, 7.30°]. Minor radius of torus-shaped ring stimulus corresponds to [0.3°, 0.31°, 0.33°, 0.34°, 0.36°, 0.37°, 0.39°, 0.40°, 0.42°, 0.44°, 0.46°, 0.48°, 0.50°, 0.52°, 0.55°, 0.57°]. It determines the size of Gabor as two times the standard deviation(*σ*) of a 2D Gaussian. The colored circle in Figure. 2A is the outline of a 2D Gaussian envelope containing 95% of the Gaussian filter (2*σ* in radius). Each ring aperture stays given eccentricity for 1.5 s (1TR) and expands or contracts in discrete steps with the same size of Gabor radius, so each ring shape stays 3 s (2TR) in the whole visual field except for the innermost (<1°) and outermost (>7.3°). Given an eccentricity, four sets of a pack of Gabor stimuli form a ring aperture (the upper panel in Fig. 2A). To minimize the spatial interaction effects between simultaneously appearing Gabor stimuli, Gabor stimuli in each set are spatially separated more than three times of the Gabor size in radius.

For the clockwise/counter-clockwise rotating wedge scan, the wedge-shaped aperture consists of two kinds of eccentricity sets (lower panel in Fig. 2A; red/magenta and blue/green). The first set consists of 7 eccentricities (1°, 1.65°, 2.39°, 3.24°, 4.22°, 5.35°, 6.64°; red/magenta colored), and the second set consists of 6 eccentricities (1.30°, 1.99°, 2.78°, 3.70°, 4.74°, 5.95°; blue/green colored). [0.30, 0.35, 0.40, 0.46, 0.52, 0.60, 0.69] or [0.32, 0.37, 0.42, 0.49, 0.56, 0.65] determines the size of Gabor stimulus as two times the standard deviation(*σ*) of a 2D Gaussian. The wedge aperture rotates in the discrete step of 11.25°. To minimize the spatial interaction effects between neighboring Gabor patches, we presented only one Gabor stimulus at a given eccentricity, composing a zigzag pattern.

The functional MRI experiment consists of 9 scans: 8 traveling scans with ring-or wedge-shaped aperture and one hemodynamic impulse response (HIRF) estimation scan. Eight traveling scans included two stimulus orientation conditions, radial and tangential orientation, four traveling apertures, CW, CCW, EXP, and CONT (CW: clockwise wedge; CCW: counter-clockwise wedge; EXP: expanding ring; CONT: contracting ring). Experiments were conducted in the following fixed order: HRF, CW_c, CW_r, EXP_r, EXP_c, CCW_c, CCW_r, CONT_r, CONT_c (_r and _c indicate radial and tangential orientation conditions, respectively). The Gabor orientation was maintained during nine cycles of each scan to obtain a robust measure of orientation-dependent fMRI time series. The fixation behavior during the experiment was assured by monitoring subjects’ performance on a fixation task, in which they had to detect any reversal in the direction of one pair of small green dots (0.07° in diameter) on a stationary red tangential circle (0.14° in diameter) at the center of the screen.

In this experimental design, we want to stress that only the stimulus orientation content was manipulated in radial or tangential orientation within a fixed spatiotemporal layout of a Gaussian envelope (e.g., CW_r vs. CW_c scan). Thus orientation-dependent changes in the anisotropy of the spatial extent of pRF at the same cortical site are due to differences in neural contribution to the pRF, not due to nuisance factors that contribute to pRF (e.g., hemodynamic response properties), because they were constant across orientation conditions.

#### MRI data preprocessing

All functional EPI images were slice-timing corrected and motion-corrected using SPM8 (Friston et al., 1996; Jenkinson et al., 2002): The measurement times for individual images were corrected by shifting the phase of all frequency components, then 3D rigid-body transformations were performed to realign image frame to the first frame of the scan. After correction, the functional data from all scans aligned to the T1-w anatomical image (reference) by application of warp matrix from the high-resolution in-plane image to the reference image (Nestares and Heeger, 2000). In-house analysis code was used in conjunction with mrTools analysis package (http://gru.stanford.edu/doku.php/mrtools). The cortical surface was constructed from a 1mm^3^ or 0.85mm^3^ T1-w image rendering of each subject using Freesurfer (Dale et al., 1999; Fischl et al., 1999) to create a flattened representation of the occipital lobe. Cortical segmentation from the inner and the outer layer of the gray matter was achieved using Jonas Larsson’s SurfRelax. This information is used in mrLoadRet to display a flattened representation of the occipital cortex. The in-house analysis codes were used in conjunction with the SPM8 and mrTools.

We manually defined the boundaries between V1, V2, and V3 visual areas on the flattened gray matter cortical surface mainly based on the meridian representation by analyzing the temporal phases of fMRI responses to rotating wedge stimuli (Engel et al., 1997).

#### Voxel selection for pRF analysis

The first 16 frames of each scan were discarded due to start-up magnetization transients. The individual voxel’s fMRI time series was converted into percent signal changes and linearly detrended to correct the slow non-physiological baseline activity components.

We applied four criteria when selecting valid V1, V2, and V3 voxels for further analysis. First, voxels showing strong response modulation with cyclic visual stimulus were selected: (1) Correlation between the measured time series and the best-fitting sinusoidal function > 0.4. (2) Voxels that responded to the HIRF stimuli (simple on-/off-stimulation of the whole visual field) with phases within ± *π*/4 around the phase of the stimulus onset. (3) Voxels with SNRs smaller than two were discarded. The signal-to-noise ratio (SNR) is computed in a Fourier domain as the amplitude of the stimulus frequency component (1/24 s) divided by the average amplitude of frequency components three times higher than the stimulus frequency for each voxel. Second, voxels that include large draining veins near the pial surface were discarded from the analysis because their fMRI signals are decoupled from neural activity both in the spatial domain (Olman et al., 2007; Polimeni et al., 2010) and in the temporal domain (Silva et al., 2000): The voxels were classified as blood-vessel-clamping if its variance was in the highest 10% of whole voxels. 55.2% ± 6.3% of total voxels (across subjects, mean ± s.d.) survived these criteria.

#### Subject-specific HIRF estimation

For HIRF estimation, 144 frames of images were acquired before the retinotopy mapping scan, and the first cycle (16 frames) was discarded due to start-up magnetization transients. The hemodynamic response was driven as the response to a briefly pulsed (3 s in duration, two times the fMRI sampling rate) full-field Gabor patches. The stimulus was repeated nine times, separated by 21-s intervals. We modeled HIRF as a difference of two gamma functions with six free parameters (Friston et al., 1998; Glover, 1999) by fitting the predicted fMRI time series to the observed time series using a least-square procedure (MATLAB’s fminsearch.m). Here, because there is HIRF variability across subjects (Aguirre et al., 1998; Handwerker et al., 2004) and the pRF estimates (especially spatial extents of pRF) depend strongly on the HIRFs (Lage-Castellanos et al., 2020; Lerma-Usabiaga et al., 2020), we derived subject specific HIRF fits. The percent variance explained is 97.68 ± 0.01% (mean ± s.d. across 29 subjects).

#### 2D elliptical pRF estimation

We used a model-based pRF analysis that was developed to estimate the stimulus-referred aggregate RF of a population of neurons within a voxel (Dumoulin and Wandell, 2008). For each voxel, pRF analysis was conducted in a two-stage procedure. The first step is a 2D isotropic Gaussian model fitting to estimate the pRF center position, and the second is a 2D elliptical Gaussian model fitting to see how stimulus orientation affects the anisotropy of pRF in the polar space.

A 2D isotropic Gaussian pRF model has three free parameters, *x*_0_, *y*_0_, *σ*, as follows: 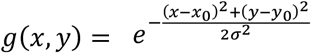. A linear overlap between the pRF, *g*(*x, y*), and a binary mask, *s*(*x, y, t*), of stimuli across time predicts the response of neural population at each measurement unit. A temporal convolution was then performed between the neural response and the subject-specific HIRF estimate, *h*(*t*0, to generate a predicted fMRI response: *y*(*t*) = *h*(*t*) * ∑_*x,y*_ *s*(*x, y, t*)*g*(*x, y*), where * denotes convolution. We use a cost function to search the best-fitting parameter set as one minus correlation coefficient between observed and predicted BOLD responses. Using MATLAB’s fminsearchbnd.m function to apply bound constraints on searching parameters with our prior knowledge. During optimizing the pRF position [x_0_, y_0_], the eccentricity was constrained to larger or equal to 0.7° and to smaller or equal to 7.9°, reflecting the visual field coverage of our stimulus (see ***visual stimulus design***). The parameter *σ* was constrained to larger or equal to .1° and to smaller or equal to 5°.

In a subsequent pRF analysis, we found the best-fitting 2D elliptical Gaussian model, which has two fixed parameters [x_0_, y_0_], and two free parameters [*σ*, aspect ratio]. The RF center was fixed with parameters determined by the 2D Gaussian model fitting results. As we are interested in the axial anisotropy of pRF along the radial and tangential axes, the elongation axis is constrained to be aligned radial or tangential axis. The aspect ratio of pRF was defined as *σ*_*radial-axis*_/*σ* _*tangential-axis*_, where *σ*_*radial-axis*_ and *σ*_*tangential-axial*_ are radial and tangential spatial extents of pRF, respectively. To generate a radially or tangentially elongated pRF model, we first get new x and y coordinates after counter-clockwise rotation angular position (*θ*), by applying 2X2 rotation matrix,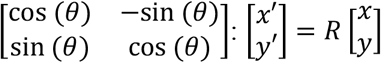. In the same manner, new coordinates of pRF position were obtained as 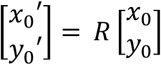. A elliptical pRF model is defined as 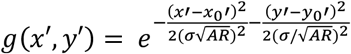.

To increase the reliability of the ellipsoid RF model fitting, we constrained the parameter space of the 2D ellipsoid Gaussian function in three ways: (1) Their pRF center position parameters are fixed with – obtained from best-fitting 2D isotropic Gaussian model. (2) The elongation axis of an ellipsoid is constrained to the radial or tangential axis. (3) The aspect ratio parameter was constrained so that the absolute log aspect ratio is smaller than 2, based on previous studies reported that voxels in early visual areas slightly elongated with an aspect ratio ranging between 1–1.67 (Merkel et al., 2018, 2020; Lerma-Usabiaga et al., 2021). Constraints with log aspect ratio < 3 do not affect our results (data are not shown).

In V1, V2, and V3, we selected valid voxels for further analysis with conservative criteria, that correlation coefficient r between observed and predicted data > 0.25 (or r > 0.3 is also tested), is satisfied all fMRI scans. In the ROIs, 28.1% ± 7.6% of selected voxels for pRF analysis (see ***Voxel selection for pRF analysis***) survived the criteria (across subject, mean ± s.d., 495.4 ± 157.1 for the number of voxels).

#### Split-half reliability of aspect ratio estimates for 29 subjects

To test the internal reliability of the aspect ratio parameter of pRF, which determines the elongated shape of pRF, we took a random sample of half (4 cycles) of the whole time series during eight cycles and ran a 2D ellipsoidal Gaussian pRF model fitting analyses between two respective split-halves. Note that because large-scale background co-fluctuations over an entire population of neurons exist whether individual neurons’ stimulus preferences match incoming visual input or not (Jack et al., 2006; Donner et al., 2008; Sirotin and Das, 2009; Choe et al., 2014; Ryu and Lee, 2018), we randomly choose half samples of time-series data. We found significant across-subject correlations between the aspect ratio parameters averaged across voxels (Pearson’s r = 0.654 or 0.719; p < e-04; voxel selection criterion correlation r > .25 or .3 was tested).

### Methods for psychophysics experiment

#### Stimuli

Stimuli were gray-scale images presented on a calibrated LCD monitor located 70 cm from subjects and set to a resolution of 1,280 × 1,024 pixels (pixel size = 0.295 × 0.293 mm). Each stimulus display consisted of a pair of targets, each positioned 3.5° from fixation and centered oppositely. The target stimuli have the same orientation of radial or tangential with 1.96 cycles-per-degree gratings, which were windowed by a two-dimensional Gaussian contrast function whose s.d. in the radial and tangential axes had a geometric average of 0.2 degrees. All stimuli were presented at 80% contrast against a uniform gray background with a mean luminance of 40 cd m−2. All stimulus properties were matched with fMRI experimental design.

#### Task

Observers participated in a circularity discrimination task (Fig. 4A; adapted from Michel et al., 2013). On each trial of the task, while fixating on the display center, the subject viewed a pair of Gabor stimuli briefly (200ms) presented at mirrored locations on the radial axis with respect to the fixation and judged which one has a shape closer to a circle. The paired stimuli always had the same orientation, either radial or tangential, but can have different contrast envelopes: one with an isotropic 2D Gaussian envelope (‘standard’) and the other with a radially or tangentially elongated ellipsoidal envelope (‘probe’). The probe’s radial-to-tangential aspect ratio (AR) was selected randomly from 9 values, which are equally spaced between 0.5 and 2 on a logarithmic scale (0.5, 0.59, 0.70, 0.84, 1, 1.19, 1.41, 1.68, 2).

All subjects first performed a practice experiment (50 trials/block, up to 8 blocks) with correct, incorrect, or late feedback. In case of incorrect, the subject can learn the task with an explanation for incorrect feedback. The subject was allowed to start the main experiment only if the task performance of the subject met the criterion with 75% correct or more. In the main experiment (100 trials/block, a total of 10 blocks), there was no feedback for each trial, and only overall performance in each block was presented. This design is to minimize the effect of understanding instruction. Participants reported which one has a shape closer to a circle by pressing the left or right arrow key when a pair of Gabor stimuli presented in horizontal or oblique meridians. When the stimuli were presented in vertical meridian, upward and downward arrow keys were used to receive their report.

#### Analysis of behavioral data: perceived anisotropy estimation by fitting psychometric curve

For each subject, we estimated the AR of the probe that appears to be a perfect circle. To compute the fractions of choosing the standard as a more circular one as a function of the AR of the probe (Fig. 4C), we assumed that decision is based on noisy perceptual estimates of the AR (Michel et al., 2013). The noise (**σ**) was proportional to the absolute value of the logarithm of perceived AR and distributed log normally. Thus, perceptual noise is the least about the Gabor stimulus with a perceived AR of 1, given that aspect ratio discrimination is best at AR=1 (Regan and Hamstra, 1992). That is,

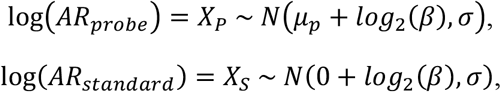

where *μ*_5_ is log(*AR*_*probe*_), and *β* is the perceived aspect ratio of the standard stimulus.

The observer’s task was to choose a stimulus with a more circular envelope. It is equivalent to determining which of the two log aspect ratios is closer to zero. In other words, the subject selects the standard whenever 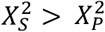 and choose the probe otherwise. We can compute the probability of choosing the standard as below:

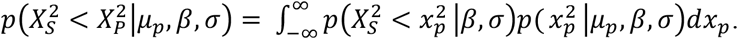

We estimated three free parameter values, [*σ, β, const*.] to maximize the sum of log-likelihoods of the observed data for each individual observer and for each stimulus orientation condition using MATLAB’s fminsearch.m.

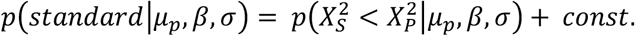

## Results

### Rationale for pRF estimation using radially and tangentially oriented gratings

To determine whether co-axial or radial bias governs the spatial representation of EVC, we inspected the spatial extents of EVC’s pRFs. To infer the bias governing the spatial representation of EVC from the spatial extents of pRFs, it is important to use appropriate stimuli for pRF estimation. For this reason, we opted to use two types of gratings differing in the orientation defined in the polar space, one oriented tangentially and the other radially (Fig. 1C). Below, we will provide the rationale for why the pRF estimation with those tangential and radial gratings can—but that with the conventional, non-oriented stimuli cannot—inform us of the bias governing the spatial representation of EVC. Because this rationale has also important implications for interpreting the results from previous studies, (which will be addressed in Discussion) it deserves a bit of elaboration.

The rationale of our experimental design is grounded in two facts. The first one is about the relationship between the organizational unit of EVC and the sampling unit of fMRI. The functional unit of EVC that is relevant to the question of our study is the orientation pinwheel, a hypercolumn within which the orientations of visual features are represented in pinwheel-like arrangement. According to the anatomical studies (Obermayer and Blasdel, 1997; Yacoub et al., 2008; Kaschube et al., 2010), the sampling unit of fMRI (2 × 2 × 2 mm^3^ voxel in our case) contains approximately 10∼30 orientation pinwheels. Thus, a single voxel’s BOLD response to an arbitrary visual input can be conceptualized as one large, virtual pinwheel’s response to that input. The second fact is the correlation between the preferred orientations and polar-angle position of single cells (Leventhal, 1983; Leventhal et al., 1984; Bauer et al., 1989; Durand et al., 2007) or local cortical sites (Freeman et al., 2011, 2013; Sun et al., 2013; Fang et al., 2022; Roth et al., 2022). From the two facts, we can deduce that the relative proportions of orientation columns within the virtual pinwheel vary systematically depending on the voxel’s polar-angle position: the preferred orientation of the locally predominant column matches the polar-angle axis of the voxel in visual space (as depicted by clusters of colored dots in Fig. 1A,B).

Critically, in the presence of this large-scale functional anisotropy in orientation representation of EVC, pRF’s spatial extent is destined to be radially elongated regardless of whether the bias governing EVC is co-axial or radial as long as pRF’s spatial extent is estimated using the conventional, non-oriented stimuli (e.g., checkerboard pattern). Suppose the governing factor is co-axial bias. Then, although orientation-nonspecific stimuli, of course, would activate all orientation columns equally, the summed orientation preferences of a voxel would be determined by the major orientation column, the preferred orientation of which is aligned to the voxel’s polar-angle position (as depicted by ellipsoids with thick contours in Fig. 1A). Thus, ironically enough, the co-axial bias in single cells’ RF or their connections leads to the radially elongated pRF anyway, given the two aforementioned facts. Needless to say, the radial bias would lead to the radially elongated pRF (Fig. 1B). This is a situation of degeneracy, where two causes bring in the same consequence, meaning that we cannot draw any conclusion about the bias governing EVC’s spatial representation form the spatial extent of pRFs estimated with the conventional, orientationally non-specific stimuli.

We avoided this degeneracy by estimating the pRFs from the BOLD responses to the gratings with radial (Fig. 1C, top) and tangential (Fig. 1C, bottom) orientations. This can be a solution to the degeneracy because we can logically decorrelate the *natural* correlation between preferred orientation and polar-angle position of visual neurons or their connections. In other words, the radial and tangential gratings would evoke the orientation-selective population within a voxel regardless of the voxel’s polar-angle position (Fig. 1D). Consequently, the co-axial and radial make distinct predictions about how the spatial extents of pRFs are elongated when pRFs are estimated with the tangentially oriented gratings: the former predicts the elongation along the tangential direction and the latter along the radial direction (Fig. 1E). Lastly, the pRFs estimated by the radial orientation will provide a reference value of spatial extent anisotropy, as follows. If co-axial bias completely governs EVC’s spatial representation, the pRFs must be elongated in the orthogonal directions but with the same degree of anisotropy between the two grating orientation conditions (Fig. 1E, left). If radial bias completely governs EVC’s spatial representation, the pRFs must be elongated in the same radial direction with the same degree of anisotropy between the two grating orientation conditions (Fig. 1E, right).

### Anisotropic spatial extent of pRF in human EVC

We acquired fMRI measurements from EVC of 29 human subjects while they viewed a pack of Gabors that were drifting slowly around a fixation spot. Since we were interested in the anisotropy in pRF spatial extent along the radial versus tangential axes in the polar space, we spatially arranged Gabors into a wedge or ring shape and made them traverse the visual field in the radial and tangential directions, respectively (Fig. 2A,B). By fitting the pRF model to the fMRI responses to the wedge and ring shapes of Gabors at a single voxel, we could estimate the voxel’s pRF location (Fig. 2C) and its spatial extents along the radial and tangential axes, respectively (Fig. 2D).

Importantly, with the rationale explained above (Fig. 1), we created the two viewing conditions in which the Gabors were oriented radially or tangentially (Fig. 2B) and estimated pRFs separately for the two conditions.To effectively estimate the radial and tangential extents of pRFs, we modeled the pRF with the 2-dimensional elliptical Gaussian function with the axes being fixed at the radial and tangential axes in the polar space. Then, for each voxel in each subject’s EVC, we fitted this model to the voxel’s fMRI time series in conjunction with the hemodynamic impulse response function (HIRF) tailored to each individual’s EVC. As a result, we estimated two pRFs for each voxel, one defined with the radially oriented patterns and the other with the tangentially oriented patterns. For each of the two pRFs, we quantified the anisotropy by contrasting two spatial extents of the pRF model, the one along the radial axis and the other along the tangential axis (*σ*_*radial-axis*_ and *σ*_*tangential-axis*_ respectively; the *σ*-defined contours of the spatial extents of pRF shown for an example voxel in Fig. 2D).

We first tested whether pRFs are anisotropic in the polar space by comparing the radial (*σ*_*radial-axis*_) and tangential (*σ*_*tangential-axis*_) spatial extents of the pRFs, separately for the two stimulus orientation conditions. In the radial orientation condition (Fig. 3A, left), the pRFs were elongated along the radial axis, significantly in all three areas of EVC (p=1.9e-05 for V1; p=0.046 for V2; p=5.8e-09 for V3; paired t-tests). Likewise, in the tangential orientation condition (Fig. 3A, right), the pRFs were elongated along the radial axis, significantly in V1 and V3 (p=0.014 for V1; p=1.5e-04 for V3; paired t-test) but not significantly in V2 (p=0.150). These results indicate that the anisotropy in the spatial extents of EVC’s pRFs is more governed by the voxels’ polar-angle position than by the stimulus orientation by showing that the pRFs are extended farther along the radial axis than along the tangential axis regardless of the stimulus orientation condition.

**Figure 3.**
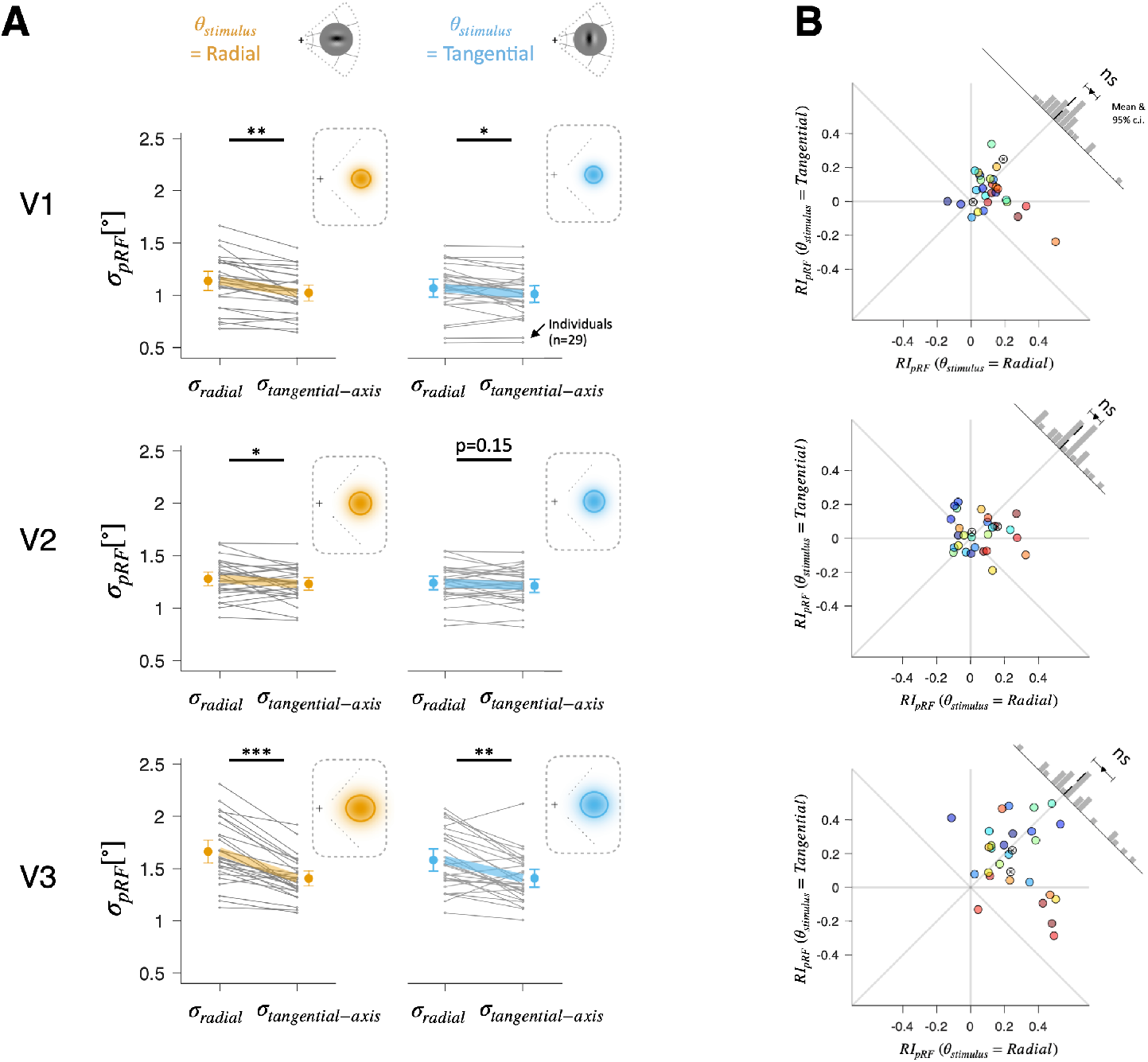
Anisotropic spatial extent of pRF in human EVC. ***A***, Comparison of the radial (*σ*_*radial-axis*_) and tangential (*σ*_*tangential-axis*_) extents of the EVC pRFs defined by the radially (yellow) and tangentially (blue) oriented gratings. Asterisks indicate the statistical significance of the comparisons (*p<0.05, **p<e-03, ***p<e-08, paired t-test) Thin gray lines represent 29 subjects. Thick colored lines and error bars represent the across-subject means and 95% confidence intervals, respectively. Insets show the across-subject averages of the spatial extents of pRF under the two stimulus orientation conditions. Here, the pRFs are centered at (3°,-3°) for illustrative purposes, and elliptical solid lines demarcate the widths of pRFs at half maximum. ***B***, Comparison in pRF’s radial elongation indices (*RI*_*pRF*_s) between the radial and tangential orientation conditions. The difference in *RI*_*pRF*_s between the two orientation conditions (upper right histogram) did not differ from 0 in all visual areas. Dots, 29 individuals; Colored dots, 27 individuals illustrated with the same color coding applied in Fig. 5 as rankings of stimulus orientation effect in perceptual anisotropy; Empty dots with ‘x’ markers inside, 2 individuals with no psychophysical data.

Next, given this predominance of radially elongated pRFs, we quantified the degree of radial elongation with *RI*_*pRF*_ by taking the signed difference between the radial and tangential extents (*RI*_5*pRF*_ = *σ*_*radial-axis*_ − *σ*_*tangential-axis*_) for further detailed inspection. The statistical tests on *RI*_*pRF*_ (paired t-test on *RI*_*pRF*_s measured during radial and tangential orientation conditions) show that the degree of radial elongation did not differ significantly between the two orientation conditions in all three areas of EVC (p=0.15 for V1; p=0.71 for V2; p=0.26 for V3; see histogram plots in upper-right panels in Fig. 3B).

We would like to explain why we opted to quantify the degree of pRF elongation based on subtractive contrast, unlike the previous fMRI studies which quantified pRF anisotropy based on divisive contrast (e.g., Merkel et al., 2018, 2020; Silva et al., 2018; Lerma-Usabiaga et al., 2021). As stated clearly in Introduction and the previous section of Results, the purpose of the current work is to determine the type of the systematic bias in *neural properties* that governs the spatial representation of EVC. With that purpose, we exploit the spatial extent of pRF to *infer* the neural bias. In this regard, we insist that the underlying bias in neural properties can be more validly captured by the subtractive contrast than the divisive contrast based on the aforementioned relationship relating the fundamental organizational unit of EVC to the sampling unit of fMRI. According to this relationship, the visual neurons that majorly contribute to the anisotropy of pRF are those whose RFs cover the edges (skirts) of visual stimuli. Furthermore, the anisotropy measure based on divisive contrast is highly dependent on the size of pRF, more exaggerating the difference as the pRF size becomes smaller. For these reasons, our anisotropy measure based on subtractive contrast is more sensitive to the bias in neural properties, on the one hand, and less invariant to the pRF size, on the other hand. Nonetheless, we note that our findings were replicated also using the anisotropy measures based on divisive contrast: there were no significant differences in the degree of radial elongation between the two stimulus orientation conditions (p=0.17 for V1; p=0.71 for V2; p=0.20 for V3; data are not shown).

### Anisotropic spatial extent estimation in human perception

The EVC hypothesis on spatial extent perception points to the high-resolution spatial representation in EVC as the neural mechanism underlying spatial extent estimation. As we reasoned in Introduction, if that is true, perceived spatial extent must be subject to the bias in neural properties—single cells’ RFs or their connections, which can be reflected in pRF of fMRI’s sampling unit—present in EVC. In the previous section, we showed that the spatial extents of EVC’s pRFs are governed by the radial bias regardless of visual input orientation, which points to the radial bias as the governing factor of EVC’s spatial representation of objects. Therefore, the EVC hypothesis predicts that the perceived spatial extents of oriented gratings must be elongated along the radial axis invariantly to the orientation of the grating.

We put this prediction to a test by measuring the anisotropy in the perceived spatial extents of the very same oriented gratings used in the fMRI experiment using a circularity-discrimination task (Regan and Hamstra, 1992; Michel et al., 2013). On each trial, subjects, while fixating at a fixation target, viewed a pair of Gabor stimuli presented at two mirrored positions on the *radial axis* from the fixation target and chose the Gabor that looks closer to the perfect circle (Fig. 4A). The paired Gabors always had the same orientation, either radial or tangential, but differing in the *physical* (i.e., true) spatial extent of contrast envelope. The *standard* Gabor had an isotropic 2D Gaussian envelope (AR=1), whereas the *probe* Gabor had an ellipsoidal envelope with a varying degree of AR (AR=[0.5 2]), where AR is the aspect ratio defined as *σ*_*radial-axis*_/*σ*_*tangential-axis*_ (Fig. 4B).

**Figure 4.**
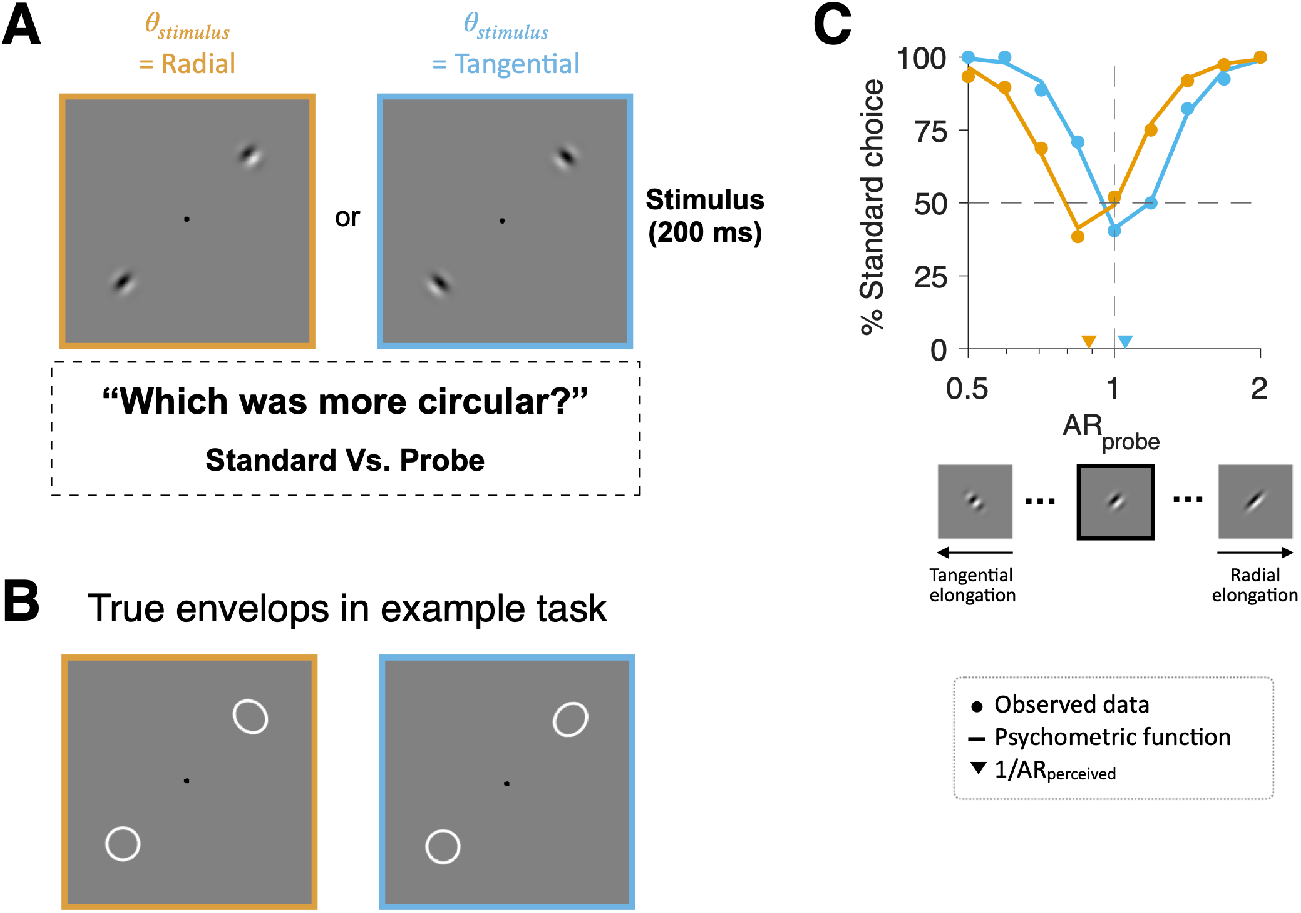
Circularity judgment task and example psychometric curves. ***A-B***, Example pairs of Gabor stimuli (***A***) and their true spatial extents (***B***) under the radial (yellow) and tangential (blue) orientation conditions. Each pair consisted of ‘standard’ and ‘probe’ stimuli of the same, radial or tangential, orientation. The aspect ratio (AR, *σ*_*radial-axis*_/*σ*_*tangential-axis*_) was fixed at 1 for the standard (lower left corners) and systematically varied for the probe (upper right corners). On each trial, subjects briefly viewed the stimuli and selected the one appearing close to a perfect circle. ***C***, Psychometric curves derived from one subject (subject 3) under the radial (yellow) and tangential (blue) orientation conditions. The fraction of choosing the standard stimulus is plotted as a function of the AR of the probe stimulus in a given trial. Note that, in the radial orientation condition (yellow curves), the observer chose the probe more frequently than the standard when the AR of the probe was 0.841. This means that the probe with AR of 0.841 looked closer to a perfect circle than the standard, whose AR is 1, indicating the co-axial elongation in perceived spatial extent. Dots, observed data; Solid lines, fitted psychometric functions; down-pointing triangle, 1/*AR*_*Percieved*_; Inset images on the x-axis, the probe Gabors at three AR values (0.5, 1, 2).

For each subject, and for each Gabor orientation condition, we derived a psychometric curve of the fraction of choosing the standard Gabor as a function of the AR of the probe Gabor (Fig. 4C), and identified the AR at which the probe Gabor appears to be a perfect circle by finding the minimum of the curve (triangles in Fig. 4C). We will refer to this physical AR defined at the minimum of the curve as *probed AR*_*physical*_. By inverting *probed A*_*physical*_, we can estimate the perceived AR of the isotropic, standard Gabor. We will refer to this perceived AR as *AR*_*perceived*_ and use its log value as the elongation index of perceived spatial extent,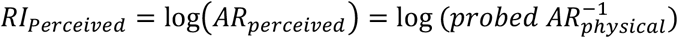.

Except for only two subjects, we could acquire the psychophysical data and estimated *RI*_*perceived*_ s from the subjects whose EVC activity was probed for the anisotropic spatial extents of pRFs. The long axis of the perceived spatial extents of the standard grating was determined by stimulus orientation (Fig. 5), elongated radially for the radially oriented Gabor (p=8.8e-06, one-sample t-test; histogram at the bottom of Fig. 5) and tangentially for the tangentially oriented Gabor (p=2.7e-04, one-sample t-test; histogram in the left of Fig. 5).These results were corroborated by the paired t-test on the difference in *RI*_*perceived*_ between the radial and tangential orientation conditions (inset histogram in Fig. 5). It should be reminded that these results were obtained from the viewing conditions where the orientations of stimuli were decorrelated with the polar-angle positions of stimuli, exactly as done in the fMRI experiment. In other words, the tangential elongation of the perceived spatial extent of the tangentially oriented Gabor should not be taken lightly because it should be considered to overcome the radial bias imposed on the grating’s spatial representation in EVC. Thus, this mismatch between the spatial extents of EVC’s pRFs and the perceived spatial extents should be taken as a serious threat to the EVC hypothesis.

**Figure 5.**
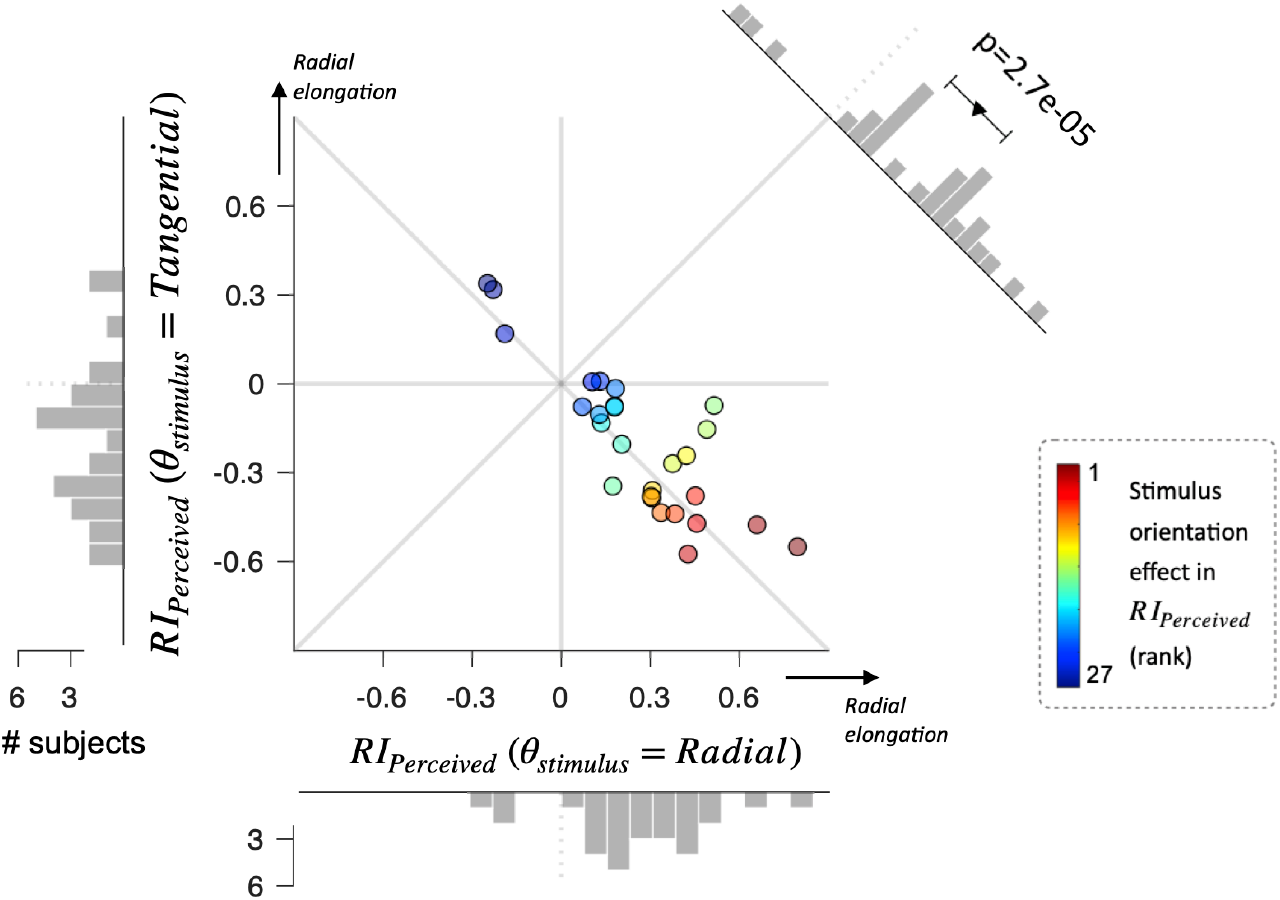
Anisotropic spatial extent estimation in human perception. The degrees of radial elongation of perceived spatial extent (*RI*_*Percieved*_) are compared between the two orientation conditions. The differences in *RI*_*pRF*_s between the two orientation conditions (upper right histogram) significantly differed from 0. Dots, 27 individuals; Colors, rank variables of stimulus orientation effect in *RI*_*Percieved*_.

### Across-subject correlation in anisotropy between EVC pRF and spatial extent estimation

The marked mismatch between the radial bias in EVC (Fig. 4) and the co-axial bias in perception (Fig. 5) strikes us as odd given the high-fidelity space representation both in EVC and in aspect ratio discrimination (Regan and Hamstra, 1992; Regan et al., 1996). One reasonable take on this seemingly odd mismatch (, on which we will elaborate in Discussion) is to take into account EVC’s location along the hierarchical streams of information processing in the primate visual cortical system. From that perspective, our results implicate that a certain mechanism existing somewhere, presumably in the downstream cortical regions, receives the high-fidelity information from EVC and transforms it such that the transformed outcomes match what is perceived. This scenario in principle can account concurrently for the high-fidelity in aspect ratio discrimination and for the mismatch in spatial bias between EVC and spatial extent perception.

Prompted by this scenario, we considered the possibility that the across-individual differences of the orientation-dependent change in the spatial extents of EVC are linearly transferred to those of perceived spatial extent. This possibility was supported by the significant correlations between the across-individual variabilities in stimulus orientation-dependent changes of *RI*_*pRF*_ and those of *RI*_*perceived*_, for all three areas of EVC (r=0.41∼0.57; p=0.001∼0.018; Fig. 6).

**Figure 6.**
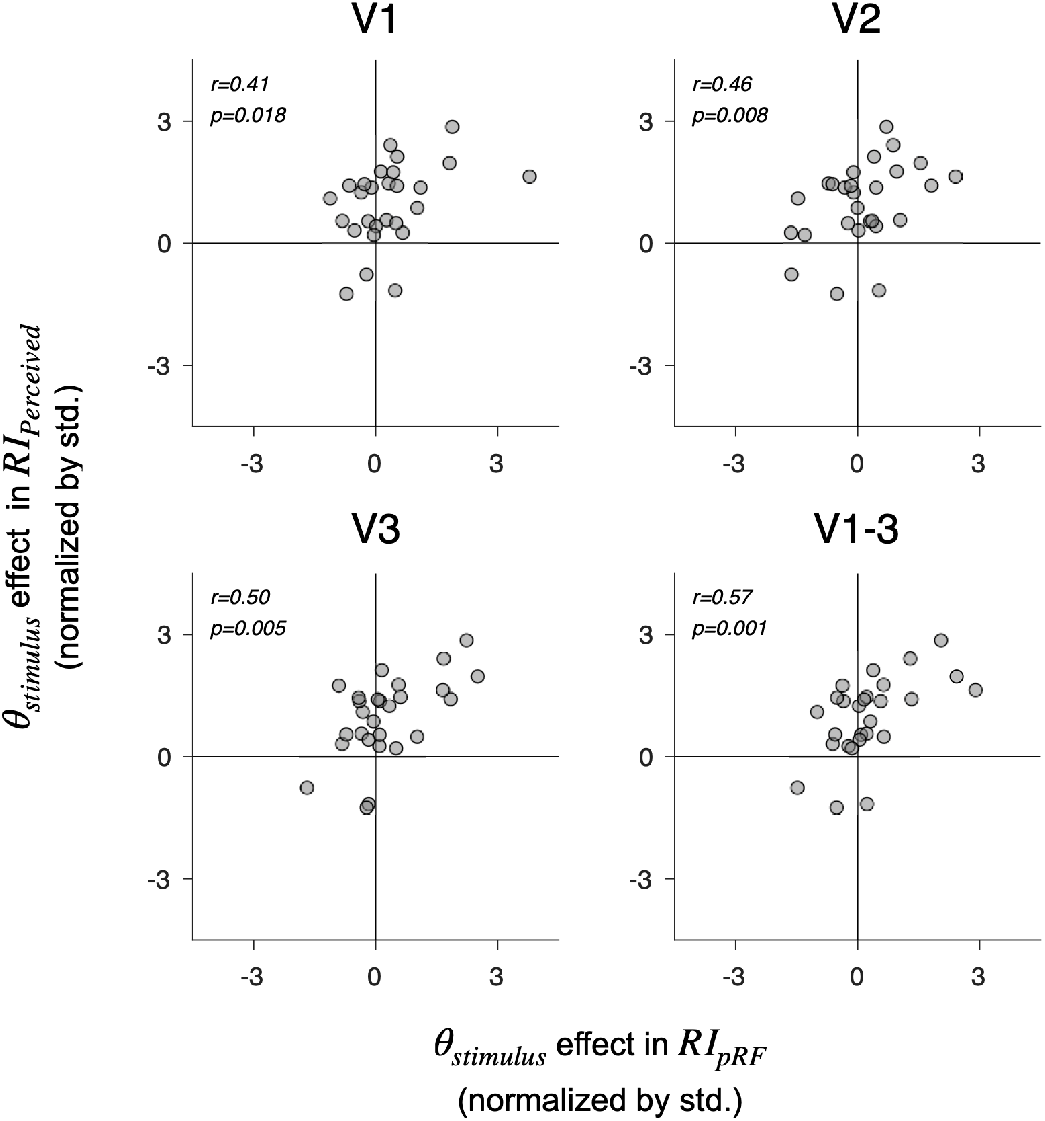
Across-individual correlation between the orientation-dependent modulation of spatial extents in EVC’s pRFs (*RI*_*pRF*_) and perceptual estimation (*RI*_*Percieved*_) for the visual areas of EVC. The modulatory effects of stimulus orientation on *RI*_*pRF*_ are plotted against those on *RI*_*Percieved*_ for individual subjects. The modulatory effects were divisively normalized by the standard deviation across subjects, so that positive values indicate that the degree of radial elongation was greater in the radial orientation condition than in the tangential orientation condition. The modulatory effects of stimulus orientation on *RI*_*pRF*_ were significantly correlated across subjects with those on *RI*_*Percieved*_. Spearman’s correlation coefficients and p-values (one-tailed test) are shown in the upper left corner of each panel.

## Discussion

Guided by an accumulated set of findings that associate spatial extent estimation with the fine spatial representation in EVC, we investigated the topographic anisotropy in EVC’s spatial representation and related it to that in spatial extent estimation. To our surprise, the radial bias pervaded the spatial extents represented by EVC, whereas the co-axial bias those perceived by humans. Despite this marked mismatch in spatial bias, the degrees to which the anisotropy of spatial extents is modulated by stimulus orientation covaried across individuals between EVC and perception. Our findings seem to require a revision of the current understanding of the biases governing EVC’s spatial representation, and about the nature of EVC’s contribution to spatial extent estimation.

### Implications of radially elongated pRFs

As we have detailed in the first Results section, the two facts must be considered when interpreting the anisotropy of pRFs: (i) the relationship between a single fMRI voxel and EVC’s orientation columns and (ii) the correlation between the preferred orientations and polar-angle positions of visual neurons or their connections. In this regard, a recent report on the radially elongated pRFs in EVC (Merkel et al., 2018) should be interpreted carefully. Since the pRFs were estimated using the conventional checkerboards, which are orientation-nonspecific, the radially elongated pRFs can arise either from the co-axial bias in conjunction with (ii) or from the genuine radial bias of horizontal connections. For that matter, the radially elongated pRFs in our study are free from such a situation of multiple interpretations since we defined the pRFs using the radially and tangentially oriented Gabors. Thus, the radially elongated pRFs in our study point to the radial bias as the governing factor of EVC’s spatial representation. Therefore, the radially elongated pRFs reported in the previous study were likely to arise from the genuine radial bias.

On a related note, the same group has recently shown that the degree of radial elongation of pRFs in EVC decreased when EVC was adapted to radially orientated stimuli (Merkel et al., 2020). The authors interpreted these results as the evidence for the co-axial bias of horizontal connections. However, adaptation-based fMRI results often allow for many alternative interpretations because the cortical adaptation can affect not just the sensitivity gain of targeted neurons but also their surround suppression interaction, or both, depending on the spatial configuration of adaptor stimuli. The full-field grating stimulus used in that study has been known to shift the orientation tunings of V1 neurons toward the adapted orientation (Kohn and Movshon, 2004; Patterson et al., 2013), which is the opposite effects the authors intended to create with adaptation.

In sum, although recent fMRI studies reported radially elongated pRFs in EVC and interpreted them as the evidence supporting the co-axial bias in horizontal connections, their findings allow for alternative interpretations, including the one in which the radial bias is a governing factor of EVC’s spatial representation.

### A fair share of the co-axial bias in EVC’s spatial representation

The win by the radial bias over the co-axial bias in our study strikes us as odd, given previous anatomical and functional studies reporting the co-axial bias. In an effort to reconcile these findings and ours, we reached a view that the presence or strength of the co-axial bias in EVC needs to be reinterpreted or reexamined with due care. Below we will explain how we reached that view.

Led by a seminal work (Bosking et al., 1997), the anatomical studies on EVC of diverse species have reported preferential horizontal connections between neurons with co-axially aligned RFs (Sincich and Blasdel, 2001; Iacaruso et al., 2017). These anatomical findings and ours are reconciled when the following aspects are considered. First, the huge differences in co-axial bias between non-primate (Bosking et al., 1997; Iacaruso et al., 2017) and primate (Sincich and Blasdel, 2001; Angelucci et al., 2002) EVCs suggest that the co-axial bias in horizontal connections might not be warranted in human EVC. This reconciliation seems plausible given the previously reported inter-species differences in other aspects of EVC’s functional architecture, including pinwheel density (Meng et al., 2012), orientation column organization (Ohki et al., 2005), and excitatory connections (Ballesteros-Yáñez et al., 2010; Chaudhuri et al., 2015; Gilman et al., 2017). Second, the presence of the co-axial bias in horizontal connections has become a matter of debate (Chavane et al., 2022). For instance, a recent study on cats’ EVC (Martin et al., 2014) could not find any co-axial alignment in the retinotopic spatial distribution of pyramidal cells’ boutons. Thus, the co-axially biased horizontal connections should not be treated as an anatomical “fact” but as an issue to be resolved.

Previous neuroimaging studies on monkey (Michel et al., 2013) and human (Park et al., 2013b; Dumoulin et al., 2014) EVC have demonstrated that EVC’s spatial representation of visual input is *modulated* by its orientation texture. These findings may seem to conflict with the predominance of radial bias over co-axial bias in our study. However, ‘co-axially elongated spatial extents’ and ‘co-axial modulation of spatial extents’ must be distinguished and do not imply each other. The former indicates that the spatial extent of an object is elongated— in an absolute sense—along the axis aligned with its orientation texture in the visual space, whereas the latter that the object’s spatial extent is *modulated* by its orientation such that it becomes more elongated—in a relative sense—along the co-orientation axis. In other words, ‘radially elongated spatial extents’ and ‘co-axial modulation of spatial extents’ can co-exist because an object’s spatial extent can remain radially elongated while its orientation modulates the degree of radial elongation. A close inspection of our results indeed hints at such a coexistence in that the degree of radial elongation was slightly attenuated in the tangential orientation condition compared to the radial orientation condition in EVC although these modulatory effects failed to reach statistical significance (Fig. 4).

In sum, the predominance of radial bias over co-axial bias found in our study does not necessarily conflict with the previous anatomical and functional studies. With this reconciliation, our findings suggest a re-evaluation of the fair share of the co-axial bias in EVC: the co-axial bias may exist, but is not as strong as previously thought, and its influence on spatial representation is modulatory in nature.

### Contribution of EVC to visual perception

The mismatch in spatial bias of EVC and perception challenges the EVC hypothesis. However, our findings should not be taken as the evidence denying EVC’s contribution to spatial extent perception. The following two aspects of our results indicate that EVC clearly contributes to spatial extent estimation but that its contribution is limited in determining the final perception of objects’ spatial extents. First, the co-axial bias in perceived spatial extents was significantly *modulated* by the polar-angle position of visual input, being more pronounced under the radial orientation condition (paired t-test, p=0.016; Fig. 5). Second, this orientation-specific modulation of the co-axial bias in perceived spatial extents was significantly anti-correlated across subjects with that of the radial bias in EVC’s pRFs (Fig. 6).

Put together, the mismatch in the direction of bias, on the one hand, and the correlation in modulatory effects, on the other hand, between EVC’s and perception’s spatial extents suggest EVC’s limited contribution to spatial extent estimation. We conjecture that EVC, at the top of the cortical stream of visual information processing, feeds its topographically refined but *radially biased* encodings of visual input into the downstream cascade, where the radially biased spatial extents are transformed into the co-axially biased spatial extents. Similar conjectures entailing the limited contribution of EVC to perception have previously been proposed as accounts of otherwise perplexing mismatches between EVC activity and visual perception during binocular rivalry (Lee et al., 2007), line drawing (Bahrami et al., 2007), visual masking (Bahrami et al., 2007), and orientation estimation (Sheehan and Serences, 2022).

It is worthy of noting the interesting correspondence between ‘the image statistics in the world versus retina’ and ‘the spatial biases in perception versus EVC’. World images abound with co-axial bias due to co-occurring edges of objects (Geisler et al., 2001; Sigman et al., 2001; Elder and Goldberg, 2002), whereas retinal images with radial bias due to saccadic eye movements (Bruce and Tsotsos, 2006; Rothkopf and Ballard, 2009; Rothkopf et al., 2009; Nandy and Tjan, 2012; Pamplona et al., 2013; Gerard-Mercier et al., 2016). From a Bayesian perspective (Hoyer and Hyvärinen, 2003; Lee and Mumford, 2003; Fiser et al., 2010), the hierarchically organized visual cortex can be understood as the architecture of inverting the process in which retinal images are generated to infer the true object in the worlds that generated the retinal images. Under this architectural scheme, the mismatch in spatial bias between EVC and perception suggests that the computational role assigned to EVC is not to represent the world images directly, but to represent the retinal images with oriented features and to hand its representation to a downstream region where the world images are recovered with the knowledge of the process of generating retinal images.

## Notes

### Competing Interest Statement

The authors have declared no competing interest.

### Summary of Updates

Title revised

